# Interruption of glucagon signaling augments islet non-alpha cell proliferation in SLC7A2- and mTOR-dependent manners

**DOI:** 10.1101/2024.08.06.606926

**Authors:** Katie C. Coate, Chunhua Dai, Ajay Singh, Jade Stanley, Brittney A. Covington, Amber Bradley, Favour Oladipupo, Yulong Gong, Scott Wisniewski, Erick Spears, Greg Poffenberger, Alexandria Bustabad, Tyler Rodgers, Nandita Dey, Leonard D. Shultz, Dale L. Greiner, Hai Yan, Alvin C. Powers, Wenbiao Chen, E. Danielle Dean

**Affiliations:** Division of Diabetes, Endocrinology and Metabolism, Department of Medicine, Vanderbilt University Medical Center, Nashville, TN; Department of Molecular Physiology and Biophysics, Vanderbilt University, Nashville, TN; The Jackson Laboratory, Bar Harbor, ME; Program in Molecular Medicine, Diabetes Center of Excellence, University of Massachusetts Chan Medical School, Worcester, MA; REMD Biotherapeutics Inc., Camarillo, CA; Department of Veterans Affairs, Tennessee Valley Healthcare System, Nashville, TN

## Abstract

**Objective:** Dysregulated glucagon secretion and inadequate functional beta cell mass are hallmark features of diabetes. While glucagon receptor (GCGR) antagonism ameliorates hyperglycemia and elicits beta cell regeneration in pre-clinical models of diabetes, it also promotes alpha and delta cell hyperplasia. We sought to investigate the mechanism by which loss of glucagon action impacts pancreatic islet non-alpha cells, and the relevance of these observations in a human islet context.

**Methods:** We used zebrafish, rodents, and transplanted human islets comprising six different models of interrupted glucagon signaling to examine their impact on delta and beta cell proliferation and mass. We also used models with global deficiency of the cationic amino acid transporter, SLC7A2, and mTORC1 inhibition via rapamycin, to determine whether amino acid-dependent nutrient sensing was required for islet non-alpha cell growth.

**Results:** Inhibition of glucagon signaling stimulated delta cell proliferation in mouse and transplanted human islets, and in mouse islets. This was rapamycin-sensitive and required SLC7A2. Likewise, *gcgr* deficiency augmented beta cell proliferation via SLC7A2- and mTORC1-dependent mechanisms in zebrafish and promoted cell cycle engagement in rodent beta cells but was insufficient to drive a significant increase in beta cell mass in mice.

**Conclusion:** Our findings demonstrate that interruption of glucagon signaling augments islet non-alpha cell proliferation in zebrafish, rodents, and transplanted human islets in a manner requiring SLC7A2 and mTORC1 activation. An increase in delta cell mass may be leveraged for future beta cell regeneration therapies relying upon delta cell reprogramming.

## 1. INTRODUCTION

Diabetes is a multifactorial disease, but insufficient insulin secretion due to inadequate functional beta cell mass and dysregulated, often increased, glucagon secretion are fundamental to essentially all forms of diabetes and contribute to hyperglycemia.^1-6^ The utility of glucagon receptor (GCGR) antagonism as a means of treating diabetic hyperglycemia has been intensively examined because it markedly improves glycemic control in diabetic rodents, non-human primates, and humans.^7-10^ More recently, studies have shown that inducible elimination of glucagon action by administration of a GCGR monoclonal antibody (GCGR-Ab) enhances beta cell survival, regeneration, and function in pre-clinical models of type 1 (T1D) and type 2 (T2D) diabetes.^11-16^

However, inhibition of glucagon action elicits adverse effects as well, including increased serum and liver lipid levels, increased blood pressure, and robust pancreatic islet alpha cell hyperplasia.^10^ We and others^17-21^ have shown that the response of islet alpha cells to GCGR antagonism is caused by disruption of the liver-alpha cell axis, which results in hyperaminoacidemia, hyperglucagonemia, and rapamycin-sensitive alpha cell proliferation through a mechanism also involving induction of a glutamine transporter, *Slc38a5*, in alpha cells.^19,20^ Recently, we extended these observations by showing that the alpha cell-enriched cationic amino acid transporter, *Slc7a2*, is required for high amino acid-stimulated mTOR activation, *Slc38a5* induction, alpha cell proliferation, and islet hormone secretion^22^, underscoring a prominent role for amino acids in the regulation of alpha cell function and phenotypes.^23^

Interestingly, pancreatic islet non-alpha cells, namely delta and beta cells, have also been shown to be impacted by GCGR antagonism.^17,24-26^ For example, constitutive ablation of the *Gcgr* gene was associated with 2- and 3-fold increases in pancreatic delta cell number and somatostatin content, respectively, and increased postnatal beta cell proliferation and mass.^17,24,25^ Furthermore, some^11,26^ but not all^18-20,27^ studies have shown that GCGR-Ab treatment increases pancreatic delta and beta cell numbers in mice and cynomolgus monkeys, respectively. However, the mechanism by which constitutive or inducible elimination of glucagon action elicits changes in islet non-alpha cells under insulin-sufficient conditions, and the relevance of these observations in a human islet context, are unknown.

Here, we used several complementary approaches to show that interruption of glucagon signaling augments islet non-alpha cell proliferation in SLC7A2- and mTORC1-dependent manners. We found that constitutive and inducible elimination of glucagon action stimulated delta cell proliferation and mass expansion in mouse and transplanted human islets, and that in mouse islets, this required mTORC1 activation and the amino acid transporter, SLC7A2.

Likewise, we found that *gcgr* deficiency increased beta cell number in a SLC7A2- and rapamycin-sensitive manner in zebrafish and promoted cell cycle engagement in rodent beta cells but was insufficient to drive a significant increase in beta cell mass in mice. Our findings highlight key differences in the regulation of beta versus delta and alpha cell proliferation and reveal a new mechanism linking inhibition of glucagon signaling to expansion of islet delta cell mass that may be leveraged for future beta cell regeneration therapies via delta cell reprogramming.^28,29^

## 2. MATERIALS AND METHODS

### 2.1. Mouse studies

All studies were performed at Vanderbilt University Medical Center and conducted in accordance with protocols and guidelines approved by the Vanderbilt University Institutional Animal Care and Use Committee. Mice were provided ad libitum access to standard rodent chow and water and housed under a 12-hour light/12-hour dark cycle. The following mice were obtained from The Jackson Laboratory: *Gcg^-/-^* (NOD.Cg-*Gcg^em1Dvs^ Prkdc^scid^ Il2rg^tm1Wjl^*/DvsJ, strain #: 029819)^30^, C57BL/6J (strain #000664), NSG (NOD.Cg-*Prkdc^scid^ Il2rg^tm1Wjl^*/SzJ, strain #: 005557)^31^, and *Slc7a2^-/-^* (B6.129S7-*Slc7a2^tm1Clm^*/LellJ, strain #: 022767).^32^ *Gcgr^-/-^* and *Gcgr^Hep-/-^*mice were generated as described previously.^17^ Wildtype (+/+) and knockout (-/-) mice obtained from heterozygous crosses were used for all experiments. For inducible elimination of glucagon action, mice were treated once per week for up to 8 weeks with control (IgG or PBS) or 10mg/kg of a humanized monoclonal antibody (10mg/kg) targeting the glucagon receptor (GCGR-Ab; “Ab-4” and REMD 2.59)^33^ via intraperitoneal (i.p.) injection.

For transplantation experiments, mouse islets were isolated by intraductal infusion of collagenase P, separated via histopaque gradient, and cultured overnight in Roswell Park Memorial Institute complete medium (RPMI; 5.6 mmol/L glucose with 10% FBS) before transplantation. Islets were isolated from 13-15-week-old *Slc7a2^+/+^* and *Slc7a2^-/-^* mice and transplanted beneath the renal capsule of contralateral kidneys in 16-18-week-old syngeneic *Slc7a2^+/+^* recipients. Following 2 weeks of engraftment, *Slc7a2^+/+^* recipient mice were injected i.p. with control IgG or GCGR-Ab (10 mg/kg) once per week for 2 weeks, after which the kidneys were harvested for tissue embedding and graft analysis as described.^22^ In the *Gcgr* mouse line, islets isolated from 14-week-old *Gcgr^+/+^*mice were transplanted beneath the renal capsule of 14-week-old syngeneic *Gcgr^Flox^* or *Gcgr^Hep-/-^* recipients as described.^17^ Human islet transplants were performed exactly as described.^19^ Human islets were obtained from the Integrated Islet

Distribution Program (https://iidp.coh.org/) or the Human Pancreas Analysis Program (https://hpap.pmacs.upenn.edu).^34,35^ Individual donor characteristics may be found in Supplemental Table 1 and Dean et al.^19^

### 2.2. Zebrafish studies

We used previously described *gcgra/b* double-knock^19^ out (abbreviated *gcgr^-/-^* here) and *slc7a2^-/-^*zebrafish lines.^22^ Proliferating beta cells were identified by incubating zebrafish embryos with 1 mmol/l 5-ethynyl-2-deoxyuridine (EdU) at four days post-fertilization (dpf) and chasing them for 24 hours. EdU was detected as described previously.^36^ Beta cell number was measured by counting *Tg(ins:H2B-mcherry)* labeled beta cells in the islet of five dpf zebrafish as described.^36^ In the rapamycin experiments, starting at 3 dpf zebrafish were treated with a concentration of 200 nM rapamycin. Treatment continued for 3 days and then beta cell number was determined at 6 dpf in *gcgr -/-* and control fish.

### 2.3. Immunofluorescence staining and image analysis

Tissue preparation and sectioning were performed as described previously.^19,22^ Tissue sections were stained for Ki67, a marker of cell proliferation (Abcam, ab15580), phosphorylated ribosomal protein S6 (pS6^240/244^), a marker of mTOR activation (Cell Signaling, #2215), and insulin (Dako, A0564), somatostatin (Santa Cruz, sc-7819), or glucagon (LSBio, LS-C202759) to mark beta, delta, and alpha cells, respectively. Alpha cells in *Gcg^-/-^* sections were identified by pro-glucagon staining (Cell Signaling, #8233). Whole pancreatic sections and islet grafts were imaged using a Scanscope FL System (Aperio Technologies) and an Olympus FV3000 laser scanning confocal microscope and analyzed using Halo image analysis software (Indica Labs). Colocalization of Ki67 with DAPI in insulin+ and somatostatin+ cells was determined by manual counting and a CytoNuclear FL v1.4 algorithm (Indica Labs). Percent beta and delta cell proliferation was quantified from at least 1500 beta and up to 1200 delta cells per animal (and at least 450 beta and 400 delta cells per donor) by dividing the number of Ki67+/insulin+ or Ki67+/somatostatin+ cells by the total number of insulin+ or somatostatin+ cells, respectively. Beta and delta cell mass was determined using an area classifier in Halo and calculated as described.^37^ Briefly, the fractional areas for insulin and somatostatin from 5-7 pancreatic sections of differing tissue depths were multiplied by the pancreas weight to obtain an estimate of total beta and delta cell mass.

### 2.4. Statistical Analyses

All data are presented as mean ± SEM. Comparisons between 2 groups were analyzed using unpaired two-tailed t tests. Comparisons between more than 2 groups were determined by one-way or two-way ANOVA with Fisher’s LSD or Tukey’s post-hoc tests. P < 0.05 denotes statistical significance. Analyses were performed using Prism 9 software.

## 3. RESULTS

### 3.1. Loss of glucagon action augments delta cell proliferation and mass expansion in mouse and transplanted human islets

To determine how interruption of glucagon signaling impacts pancreatic islet non-alpha cells, we first measured delta cell proliferation and mass in two different mouse models: one with constitutive global glucagon deficiency (*Gcg^-/-^*) and another with inducible elimination of glucagon action via treatment with a GCGR-Ab. We found that delta cell proliferation, quantified as the percentage of Ki67-positive delta cells, was increased by 4.7- and 6.2-fold in *Gcg^-/-^* and GCGR-Ab treated mice, respectively, compared with controls (**Fig 1A-C**). This resulted in a ∼ 3-fold increase in delta cell mass compared to controls (**Fig 1D-F**). Consistent with earlier reports^24,25^, we also observed a distinctive shift in the distribution of somatostatin-positive cells from the mantle to the core of the islet in both *Gcg^-/-^*and GCGR-Ab treated mice (**Fig 1A and 1D**). To evaluate the translational relevance of these findings, we measured delta cell proliferation in human islets transplanted into immunocompromised recipient mice (i.e., NSG) treated with IgG (control) or GCGR-Ab and found a 3.5-fold increase in the percentage of Ki67-positive delta cells after 4 weeks of treatment (**Fig 1G-I**). Our findings suggest that interruption of glucagon signaling promotes islet delta cell mass expansion by stimulating delta cell proliferation in mouse and transplanted human islets.

**Figure 1.**
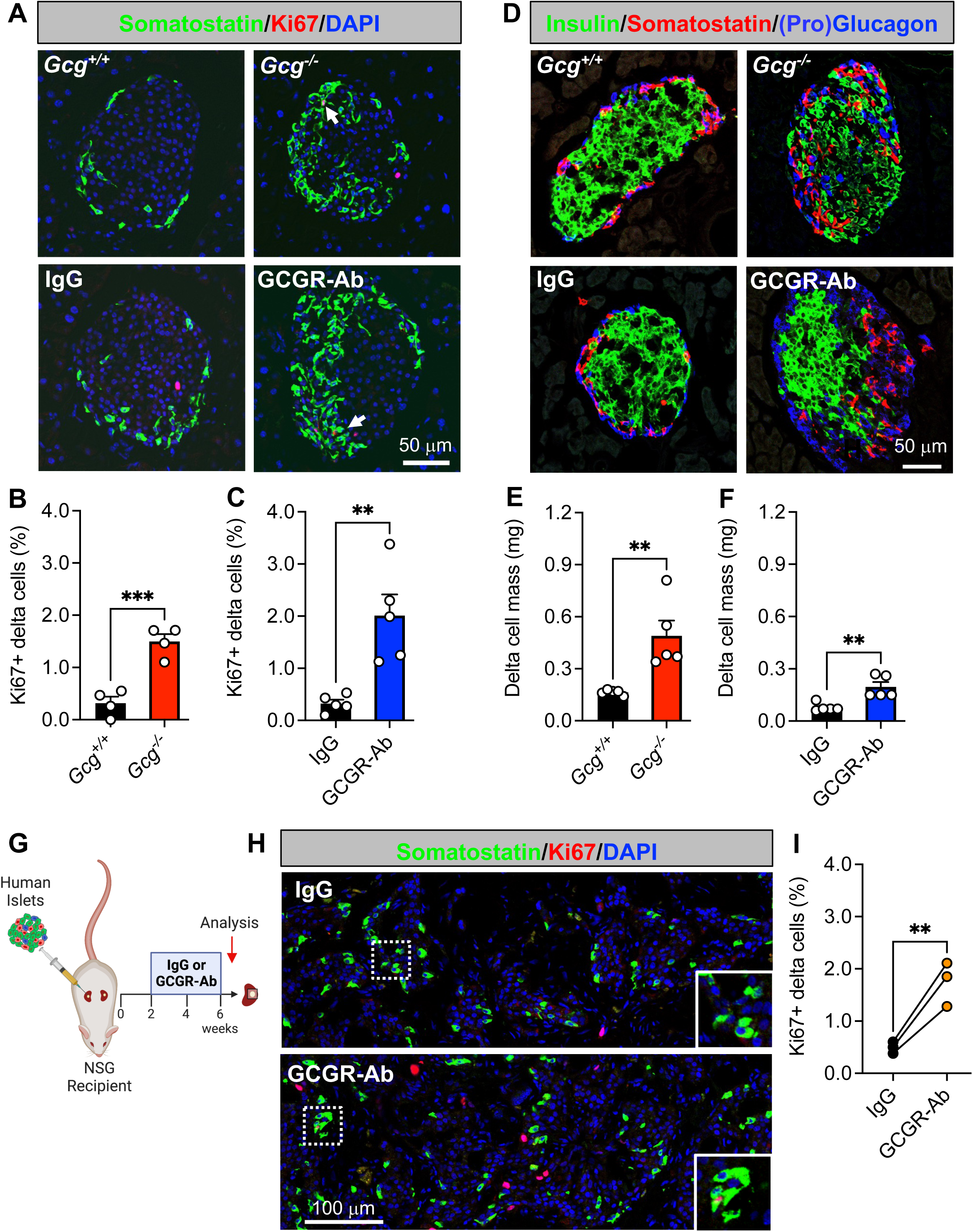
Loss of glucagon action augments delta cell proliferation and mass expansion in mouse and transplanted human islets. (**A**) Representative images of pancreatic islet and delta cell proliferation in Gcg^+/+^/Gcg^-/-^ (upper row) and IgG/GCGR-Ab-treated C57BL6 (bottom row) mice. Somatostatin (green), Ki67 (red), and DAPI (blue) are shown. White arrows indicate Ki67+ somatostatin+ cells. (**B-C**) Quantification of pancreatic islet delta cell proliferation in (**B**) Gcg^+/+^ (black bar) and Gcg^-/-^ (red bar) mice (n=1-2 females and 2-3 males per genotype) and (**C**) control IgG (black bar) and GCGR-Ab-treated (blue bar) mice (all males, unpaired t test, ***p < 0.001 versus Gcg^+/+^, **p < 0.01 versus IgG). (**D**) Representative images of pancreatic islet hormones in Gcg^+/+^/Gcg^-/-^ (upper row) and IgG/GCGR-Ab-treated (bottom row) mice. Insulin (green), somatostatin (red), and pro-glucagon (Gcg^+/+^/Gcg^-/-^; blue) or glucagon (IgG/GCGR-Ab; blue) are shown. (**E-F**) Pancreatic islet delta cell mass in (**E**) Gcg^+/+^ (black bar) and Gcg^-/-^ (red bar) mice (n=1-2 females and 3 males per genotype) and (**F**) control IgG (black bar) and GCGR-Ab-treated (blue bar) mice (all males, unpaired t test, **p < 0.01 versus Gcg^+/+^ or IgG). (**G**) Schematic of approach for human islet subcapsular renal transplantation in NSG recipient mice followed by control IgG or GCGR-Ab treatment. Created with BioRender.com (**H**) Representative images of delta cell proliferation in human islet grafts after 4 weeks of control IgG (upper row) or GCGR-Ab-treatment (bottom row). Grafts were immunostained for somatostatin (green), Ki67 (red), and DAPI (blue). White dashed boxes indicate regions selected for insets. (**I**) Quantification of delta cell proliferation in transplanted human islets in control IgG (black circles) and GCGR-Ab-treated (orange circles) mice (n=3 donors [see Supplemental Table 1], unpaired t test, **p < 0.01 versus IgG).

### 3.2. SLC7A2 and mTORC1 activation are required for delta cell proliferation in response to interrupted glucagon signaling

To gain insight into the mechanism(s) that may be driving delta cell proliferation in *Gcg^-/-^* and GCGR-Ab treated mice, we first measured delta cell proliferation in mice with constitutive global inactivation of the cationic amino acid transporter, SLC7A2. We and others^18-20^ have shown that GCGR-Ab-induced hyperaminoacidemia, especially arginine and glutamine, triggers pancreatic islet alpha cell proliferation in an mTOR-dependent manner, and that this response requires SLC7A2.^22^ Here, we found that SLC7A2 is also required, at least in part, for delta cell proliferation in response to GCGR-Ab treatment since the percentage of Ki67-positive delta cells was abrogated, albeit incompletely, in *Slc7a2* knockout (-/-) mice compared with wild-type (+/+) controls (**Fig 2A-B**). To delineate the intracellular signaling pathway involved, we immunostained for phosphorylated (p) S6 protein, a downstream target of mTOR kinase, and observed an 18-fold increase in the percentage of pS6-positive delta cells in islets of *Gcg^-/-^* compared with *Gcg*^+/+^ mice, indicative of mTOR activation (**Fig 2C-D**). To determine whether mTOR signaling was required for delta cell proliferation, we co-treated mice with GCGR-Ab and rapamycin (RAPA), an mTOR inhibitor, and found that RAPA abolished GCGR-Ab-induced delta cell proliferation in C57BL6 mice (**Fig 2E-F**). These data indicate that loss of glucagon action stimulates RAPA-sensitive delta cell proliferation through a mechanism requiring, at least in part, the amino acid transporter SLC7A2.

**Figure 2.**
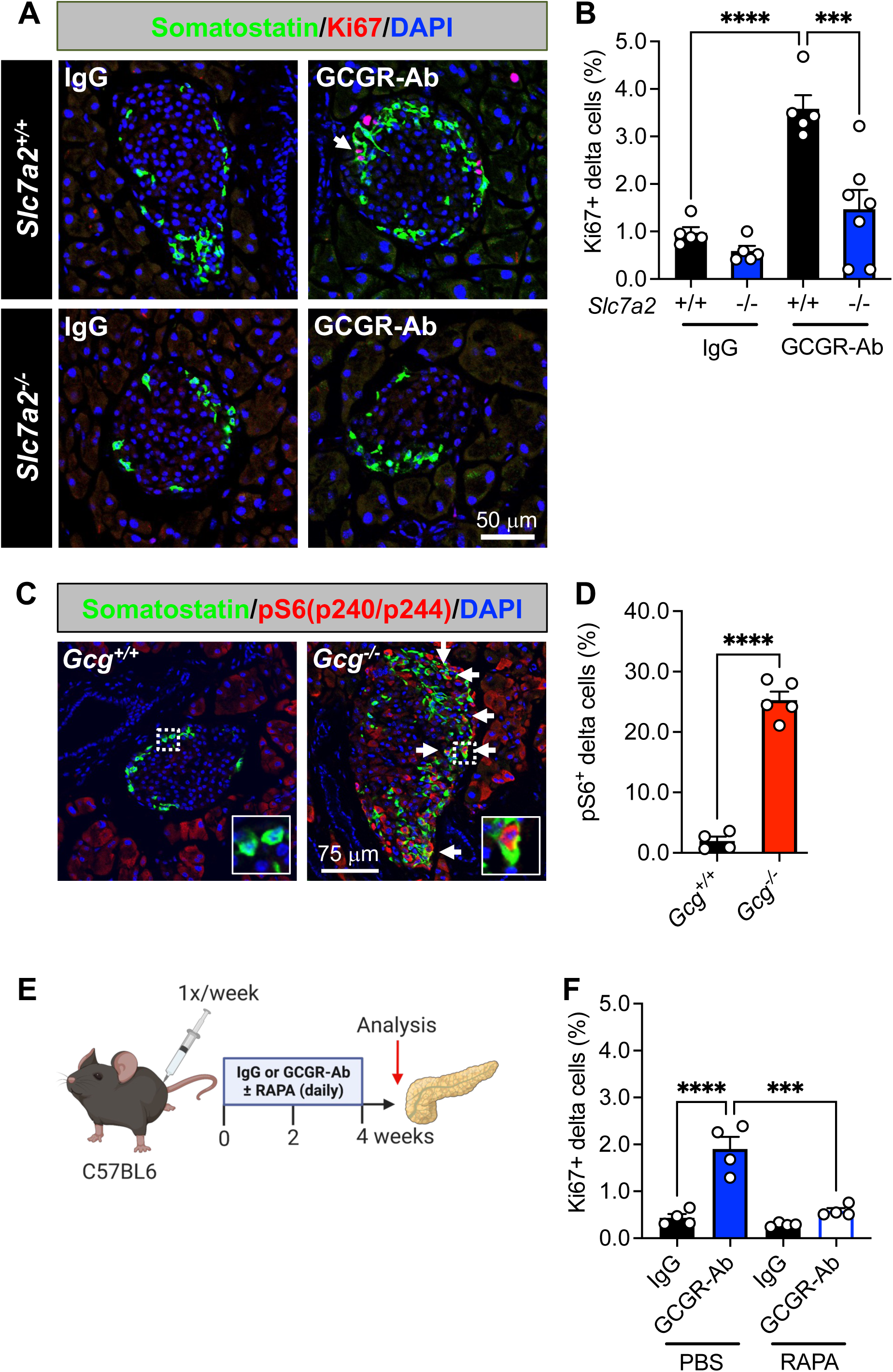
SLC7A2 and mTOR activation are required for delta cell proliferation in response to interrupted glucagon signaling. (**A**) Representative images of pancreatic islet delta cell proliferation in *Slc7a2^+/+^* (upper row) and *Slc7a2^-/-^* (bottom row) IgG/GCGR-Ab-treated mice. Somatostatin (green), Ki67 (red), and DAPI (blue) are shown. White arrows indicate Ki67+ somatostatin+ cells. (**B**) Quantification of pancreatic islet delta cell proliferation in *Slc7a2^+/+^* (black bars) and *Slc7a2^-/-^*(blue bars) IgG or GCGR-Ab-treated mice (n=4 females and 1-3 males per genotype, one-way ANOVA with Tukey’s multiple comparisons test, ****p < 0.0001 versus *Slc7a2^+/+^* IgG, ***p < 0.001 versus *Slc7a2^+/+^* GCGR-Ab). (**C**) Representative images of pancreatic islets in *Gcg^+/+^*/*Gcg^-/-^*mice immunostained for somatostatin (green), phosphorylated ribosomal protein S6 (pS6^240/244^; red), and DAPI (blue). White arrows indicate pS6+ somatostatin+ cells. White dashed boxes indicate regions selected for insets. (**D**) Quantification of the percentage of pS6+ somatostatin+ cells in *Gcg^+/+^* (black bar) and *Gcg^-/-^* (red bar) pancreatic islets (n=1-2 females and 3-4 males per genotype, unpaired t test, ****p < 0.0001 versus *Gcg^+/+^*). (**E**) Schematic of approach for IgG/GCGR-Ab (once weekly) and rapamycin (RAPA; once daily) co-treatment in C57BL6 mice. Created with BioRender.com (**F**) Quantification of pancreatic islet delta cell proliferation in mice co-treated with IgG (black bars) or GCGR-Ab (blue and white bar) and PBS or RAPA (all males, one-way ANOVA with Tukey’s multiple comparisons test, ****p < 0.0001 versus IgG, ***p < 0.001 versus PBS/GCGR-Ab).

### 3.3. Loss of glucagon receptor function stimulates beta cell proliferation in a species-specific manner

Since the response of islet delta cells to interrupted glucagon signaling resembled that of alpha cells shown previously^17-20,22,24,25,36,38^ we wanted to determine if this mechanism was likewise conserved in beta cells under insulin-sufficient conditions. First, we examined a zebrafish model with constitutive global deficiency of both forms of the *gcgr^36^* (*gcgra/b^-/-^*, abbreviated *gcgr^-/-^*) and found that beta cell proliferation and number were both increased in *gcgr^-/-^* zebrafish compared to WT (*gcgr^+/+^*) controls (**Fig 3A-B**). Furthermore, this response was significantly blunted in *slc7a2^-/-^;gcgr^-/-^*double mutants, or in *gcgr^-/-^* zebrafish treated with RAPA (**Fig 3C-D**). These data indicate that the mechanism for GCGR deficiency-induced delta and alpha cell mass expansion is conserved in zebrafish beta cells.

**Figure 3.**
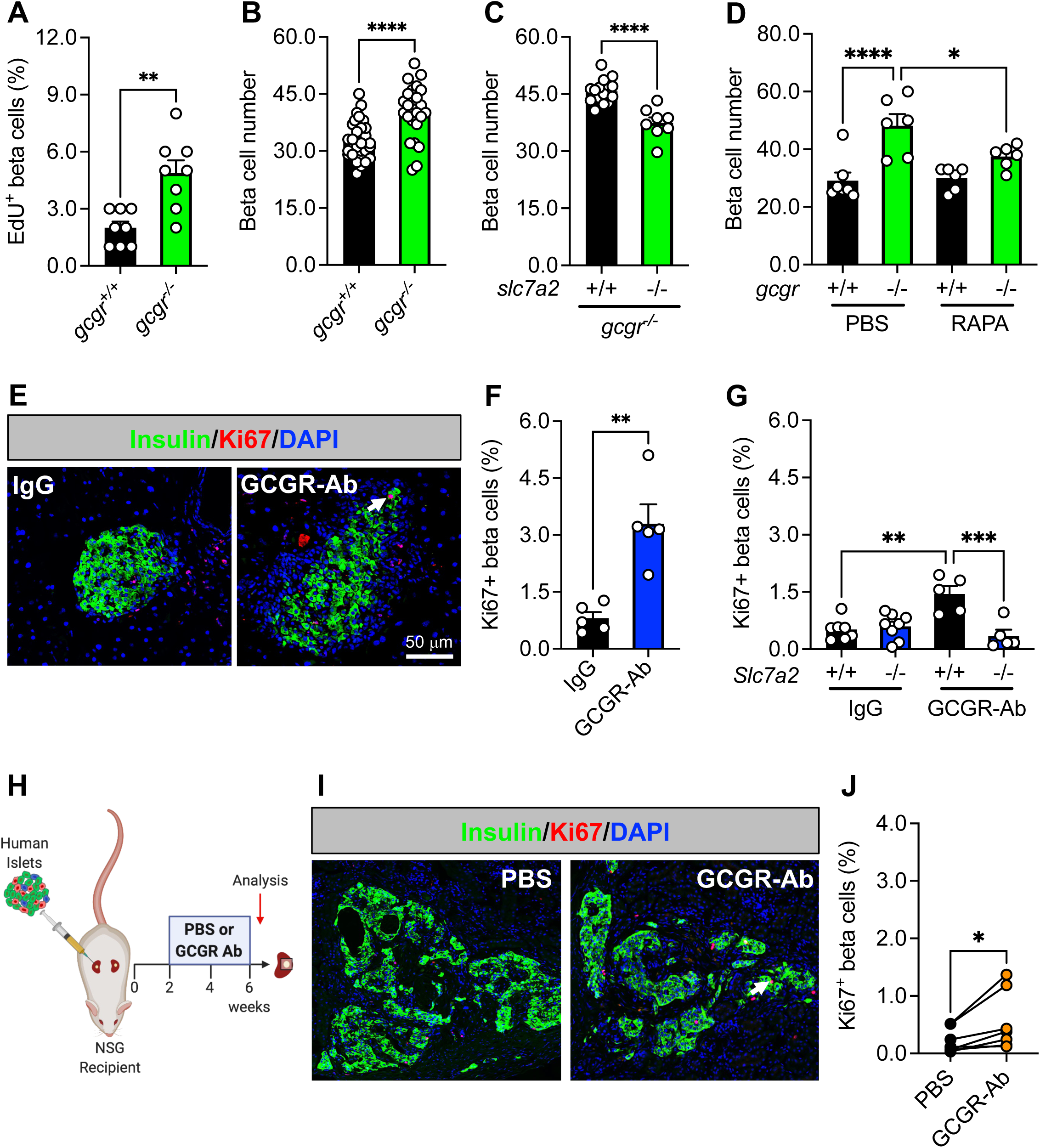
Loss of glucagon receptor function stimulates beta cell proliferation in a species-specific manner. (**A**) Beta cells stained for EdU to assess their proliferation in 5 dpf (days post-fertilization) wild-type (*gcgr^+/+^*, black bar) and *gcgra/b^-/-^*(abbreviated *gcgr^-/-^*, green bar) zebrafish (n=8 per group, unpaired t test, **p < 0.01 versus *gcgr^+/+^*). (**B**) Beta cell number in 5 dpf *gcgr^+/+^* (black bar) and *gcgr^-/-^* (green bar) zebrafish (n=24-30 per group, unpaired t test, ****p < 0.0001 versus *gcgr^+/+^*). (**C**) Beta cell number after knockdown of *slc7a2* (+/+, black bar; -/-, green bar) in 5 dpf *gcgr^-/-^* zebrafish (n=8-15 per group, unpaired t test, ****p < 0.0001 versus *slc7a2^+/+^*). (**D**) Beta cell number in 6 dpf g*cgr^+/+^* (black bar) and *gcgr^-/-^* (green bar) zebrafish after 3 days of treatment with PBS or RAPA (n=6-7 per group, one-way ANOVA with Fisher’s LSD, ****p < 0.0001 versus PBS/*gcgr^+/+^*, * p < 0.05 versus PBS/*gcgr^-/-^*). (**E**) Representative images of pancreatic islet beta cell proliferation in IgG/GCGR-Ab-treated C57BL6 mice. Insulin (green), Ki67 (red), and DAPI (blue) are shown. White arrow indicates a Ki67+ insulin+ cell. (**F**) Quantification of pancreatic islet beta cell proliferation in control IgG (black bar) and GCGR-Ab-treated (blue bar) mice (all males, unpaired t test, **p < 0.01 versus IgG). (**G**) Quantification of pancreatic islet beta cell proliferation in *Slc7a2^+/+^* (black bars) and *Slc7a2^-/-^* (blue bars) IgG or GCGR-Ab-treated mice (n=2-5 females and 3-6 males per group, one way ANOVA with Tukey’s multiple comparisons test, **p < 0.0001 versus *Slc7a2^+/+^* IgG, ***p < 0.001 versus *Slc7a2^+/+^*GCGR-Ab). (**H**) Schematic of approach for human islet subcapsular renal transplantation in NSG recipient mice followed by PBS or GCGR-Ab treatment. Created with BioRender.com (**I**) Representative images of beta cell proliferation in human islet grafts after 4 weeks of PBS or GCGR-Ab-treatment. Grafts were immunostained for insulin (green), Ki67 (red), and DAPI (blue). Dashed yellow lines indicate kidney-graft boundary. (**J**) Quantification of beta cell proliferation in transplanted human islets in PBS (black circles) or GCGR-Ab-treated (orange circles) mice (n=2 female and 5 male donors [see Supplemental Table 1], unpaired t test, *p < 0.05 versus PBS).

To examine the response of beta cells in a mammalian context, we treated adult C57BL6 mice with IgG or GCGR-Ab for 8 weeks and observed a 4.1-fold increase in the percentage of Ki67 positive beta cells (**Fig 3E-F**). This measure of beta cell proliferation was also increased in mice with constitutive global inactivation of glucagon signaling (**Fig S1A-B**), as shown previously.^25^ Furthermore, we detected an increase in beta cell proliferation in *Gcgr^+/+^*islets only when transplanted into liver-specific *Gcgr^-/-^* (*Gcgr^Hep-/-^*) and not *Gcgr^Flox^* mice (**Fig S1C-E**), supporting the premise that inhibition of hepatic glucagon action produces circulating factors that augment islet cell proliferation.^15,18-20^

To determine whether the amino acid transporter SLC7A2 was required for beta cell proliferation, we treated *Slc7a2^+/+^* and *Slc7a2^-/-^* mice with IgG or GCGR-Ab for 2 weeks and found that the GCGR-Ab-induced increase in Ki67-positive beta cells (∼2.8-fold) was abolished in *Slc7a2^-/-^* islets (**Fig 3G**), and in *Slc7a2^-/-^*, but not *Slc7a2^+/+^*, islets transplanted into *Slc7a2^+/+^* recipients (**Fig S2A-C**). These data suggest that SLC7A2 is required in an islet autonomous manner for GCGR-Ab-induced beta cell proliferation. In contrast to the response of mouse delta and zebrafish beta cells, however, we did not detect a coordinate increase in beta cell mass in GCGR-Ab-treated mice (**Fig S2D**). The discordance between beta cell proliferation and mass did not appear to be due to a detectable increase in beta cell death since we did not detect any TUNEL positive beta cells in islets of GCGR-Ab treated mice (data not shown). These findings imply that inducible elimination of glucagon action is sufficient to stimulate beta cell cycle entry, but insufficient to promote cell cycle completion in healthy mice.

Lastly, we sought to evaluate the translational relevance of these observations by measuring beta cell proliferation in transplanted human islets and likewise observed an increase in the mean percentage of Ki67-positive beta cells following 4 weeks of GCGR-Ab treatment (**Fig 3H-J**), though the magnitude of the response varied widely among donors and was lower than that of human delta cells.

## 4. DISCUSSION

The mechanism linking inhibition of glucagon signaling to changes in pancreatic islet delta and beta cells, and the relevance of these observations in a human islet context, are unknown. In this study, we used zebrafish, rodents, and transplanted human islets comprising six different models of altered glucagon signaling to show that inhibition of glucagon action stimulated delta and beta cell proliferation via SLC7A2- and mTORC1-dependent mechanisms. Constitutive global deletion of *Gcg*, or inducible elimination of glucagon signaling with a GCGR-Ab, promoted RAPA-sensitive delta cell proliferation and mass expansion through a mechanism requiring the cationic amino acid transporter, SLC7A2. Likewise, we identified an increase in beta cell proliferation and mass in *gcgr-*deficient zebrafish that was abrogated upon deletion of SLC7A2 or treatment with RAPA. While constitutive and inducible inhibition of glucagon signaling also augmented the percentage of Ki67-positive beta cells in rodent islets, this resulted in only a modest, non-significant increase in beta cell mass. Consistent with previous reports in human alpha cells^19,20^, we also showed that GCGR-Ab treatment stimulated human delta, and to a lesser extent beta, cell proliferation in transplanted islets, highlighting the translational relevance of our observations.

We and others^18-20^ showed previously that GCGR antagonism elicits hyperaminoacidemia, which promotes mTOR-dependent alpha cell hyperplasia through a mechanism involving induction of SLC38A5, a glutamine transporter, in alpha cells.^19,20^ More recently, we found that SLC7A2, an arginine transporter, is the most highly expressed amino acid transporter in zebrafish, rodent, and human alpha cells and required for activation of mTORC1 signaling, induction of SLC38A5, and stimulation of alpha cell proliferation in response to GCGR antagonism.^22^ Here, we extended these observations by demonstrating that SLC7A2 and mTORC1 activation are likewise required for islet non-alpha cell proliferation in response to interrupted glucagon signaling. This was surprising since the expression of *Slc7a2* is much lower in delta and beta cells compared to alpha cells.^22^ Because we used mice with constitutive global deletion of SLC7A2, we cannot delineate the direct versus indirect requirement for this transporter on delta and/or beta cell proliferation. Notwithstanding, we showed that GCGR-Ab treatment augmented the percentage of Ki67-positive beta cells in transplanted donor islets from *Slc7a2^+/+^*, but not *Slc7a2^-/-^*, mice supporting at least an islet-autonomous role for this transporter in amino acid-regulated cell proliferation. However, the possibility cannot be excluded that GCGR antagonism-dependent proliferative signal(s) emanating from SLC7A2-enriched alpha cells act in a paracrine manner on neighboring delta and/or beta cells to promote their proliferation. Future studies using conditional *Slc7a2* gene targeting approaches in islet delta, beta, and/or alpha cells will be necessary to delineate its requirement in specific endocrine cell subsets.

We were also surprised to find that mTORC1 signaling was required for islet non-alpha cell proliferation, since previous studies reported GCGR-Ab-mediated activation of S6 protein, a downstream target of mTORC1, in only rare beta^20^ or delta^19^ cells, and the results are conflicting. Here, we observed an 18-fold increase in the percentage of pS6-positive delta cells in *Gcg^-/-^* islets, indicative of heightened mTORC1 activity. It is possible that the pattern of S6 activation in islet cells differs between models of constitutive versus inducible elimination of glucagon action. Nevertheless, we showed that pharmacologic inhibition of mTORC1 signaling with RAPA abolished delta cell proliferation in GCGR-Ab-treated mice, and beta cell proliferation in *gcgr* deficient zebrafish, supporting a central role for mTORC1-dependent nutrient sensing in islet non-alpha cell proliferation in non-diabetic models.

Our results agree with previous studies in *Gcgr^-/-^* mice, which showed up to 3.5-fold increases in islet delta cell number, mass, and pancreatic somatostatin content.^24,25^ These studies also identified a shift in the labeling pattern of somatostatin-positive cells from being restricted to the mantle zone in *Gcgr*^+/+^ islets to being scattered within the core as well in *Gcgr^-/-^* islets, much like alpha cells.^24,25^ We confirmed and extended these observations in *Gcg^-/-^* mice by showing that the increase in delta cell mass was due, at least in part, to delta cell proliferation as evidenced by a 4.7-fold increase in the percentage of Ki67-positive delta cells. More recently, Gu et al.^26^ showed that treatment of C57BL6 mice with a GCGR-Ab at a weekly dose of 5 mg/kg for 4 weeks increased islet delta cell number by ∼35% in association with marginally significant (P=0.05) delta cell proliferation. Conversely, we showed that treatment of C57BL6 mice with a GCGR-Ab at a weekly dose of 10 mg/kg for 8 weeks triggered a 6.2-fold increase in delta cell proliferation and a 2.5-fold increase in delta cell mass. These discrepancies are likely explained by differences in the dose and/or duration of GCGR-Ab exposure, since Kim et al.^20^ also found no change in pancreatic delta cell mass in C57BL6 mice after 3 weeks of GCGR-Ab treatment at a weekly dose of only 3 mg/kg.

The impact of interrupted glucagon signaling on beta cell proliferation and mass expansion under insulin-sufficient conditions is inconclusive. Several studies^18-20,27,30^, primarily in rodent models of inducible elimination of glucagon action, have failed to detect an increase in beta cell proliferation or mass. On the other hand, constitutive global or liver-specific ablation of the *Gcgr* gene was associated with increased postnatal beta cell proliferation^25^ and up to a 1.7-fold increase in beta cell mass.^17,25^ Furthermore, Xi et al.^11^ found a ∼20% increase in the percentage of insulin-positive cells in islets of healthy cynomolgus monkeys treated with GCGR-Ab at a weekly dose of 60 mg/kg for 13 weeks, but it is unknown if this was due to an increase in beta cell proliferation. Here, we showed a significant increase in beta cell mass in *gcgr*-deficient zebrafish, but not in GCGR-Ab treated mice, despite cell cycle engagement as evidenced by 2- to 4-fold increases in the percentage of Ki67 positive beta cells. An uncoupling of beta cell cycle entry from cell cycle completion was also observed by Furth-Lavi and colleagues^39^ in a diabetic rodent model of extreme beta cell-ablation, where some beta cells were capable of entering the cell cycle but failed to complete it under conditions of severe, but not moderate, hyperglycemia. In pre-clinical models of T1D and T2D, however, GCGR antagonism improves glycemia and readily promotes the regeneration of functional beta cell mass through, for example, beta cell proliferation and alpha-to-beta-cell transdifferentiation.^11-16^ These studies suggest that the blood glucose level and/or magnitude of insulin deficiency may be determinants of beta cell regenerative capacity, and by extension, beta cell mass expansion, in diabetic rodent models. We posit that under healthy, insulin-sufficient conditions, rodent beta cells, but not alpha or delta cells, require additional proliferative signals – or disinhibition of repressive signals – for GCGR antagonism to promote cell cycle completion and an increase in beta cell mass. Tight control over insulin production is necessary to protect against hypoglycemia and may reflect physiologic autoregulation of beta cell mass in healthy animals. Species-specific differences in the response of beta cells to GCGR loss likely reflect the heightened plasticity, regenerative capacity and developmental stages of zebrafish versus mammalian islet cells.^40-43^ Future studies aimed at identifying the extra- and/or intra-cellular effectors that couple beta cell cycle engagement with cell cycle completion in health and diabetes will shed new light on unique mechanisms of mammalian beta cell regulation.

These studies demonstrate a physiologic role for glucagon in the regulation of islet delta and beta cell mass and expose notable differences in their proliferative capacity in healthy mice. Future studies should address how amino acid-dependent nutrient sensing stimulates islet non-alpha cell proliferation, and whether GCGR-Ab induces delta cell mass expansion in diabetic models. By enhancing our understanding of the mechanism(s) linking inhibition of glucagon action to expansion of islet cell mass, we may improve our ability to mitigate the negative side effects of GCGR antagonism while leveraging its favorable effects, including the possibility for beta cell regeneration therapies relying upon delta (and/or alpha) cell reprogramming in T1D and T2D.^26,28,29^

## DECLARATION OF COMPETING INTERESTS

The authors declare no conflicts of interests.

## CRediT AUTHORSHIP CONTRIBUTION STATEMENT

**Katie C. Coate:** Conceptualization, Formal analysis, Validation, Investigation, Visualization, Supervision, Project Administration, Funding acquisition, Writing – original draft, Writing – review and editing; **Chunhua Dai:** Conceptualization, Methodology, Formal analysis, Validation, Investigation, Visualization, Supervision, Project Administration, Writing – review and editing; **Ajay Singh:** Investigation, Validation, Formal analysis; **Jade Stanley:** Investigation, Validation, Formal analysis; **Brittney A. Covington:** Investigation, Validation, Formal analysis; **Amber Bradley:** Software, Investigation, Validation; **Favour Oladipupo:** Investigation, Validation, Formal analysis; **Yulong Gong:** Investigation, Validation, Formal analysis; **Scott Wisniewski:** Investigation, Validation, Formal analysis; **Erick Spears:** Resources, Methodology, Investigation; **Greg Poffengerber:** Methodology; **Alexandra Bustabad:** Investigation, Validation, Formal analysis; **Tyler Rodgers:** Investigation, Validation, Formal analysis; **Nandita Dey:** Investigation, Validation; **Leonard D. Shultz:** Resources, Writing – review and editing; **Dale L. Greiner:** Resources, Writing – review and editing; **Hai Yan:** Resources; **Alvin C. Powers:** Conceptualization, Resources, Writing – review and editing, Supervision, Project administration, Funding Acquisition; **Wenbiao Chen:** Conceptualization, Resources, Writing – review and editing, Supervision, Project administration, Funding Acquisition; **E. Danielle Dean:** Conceptualization, Methodology, Validation, Investigation, Formal Analysis, Visualization, Resources, Writing – review and editing, Supervision, Project administration, Funding Acquisition

## ACKNOWLEDGEMENTS

This research was performed using resources and/or funding provided by the following: JDRF SRA-149-Q-R (to A.C.P. and E.D.D.), R01DK117147 (to W.C. and A.C.P.), an administrative supplement to R01DK117147 (to K.C.C.), R01DK132669 (to E.D.D.), K01DK117969 (to E.D.D.), the Human Islet Research Network (UC4 DK104211 and DK112232), R24DK106755 (to A.C.P.), DK104218 and OD0426640 (to D.L.G and L.D.S.), P30DK020593 (Vanderbilt Diabetes Research and Training Center), P30DK058404 (Vanderbilt University Medical Center’s Digestive Diseases Research Center), T35DK007383 (Vanderbilt Student Research Training Program), and the Department of Veterans Affairs (BX000666 to A.C.P.). Islet isolation and slide scanning was performed using the Islet and Pancreas Analysis (IPA) Core supported by the Vanderbilt Diabetes Research Center (NIH grant P30DK020593).

---

**Supplemental Figure 1.**
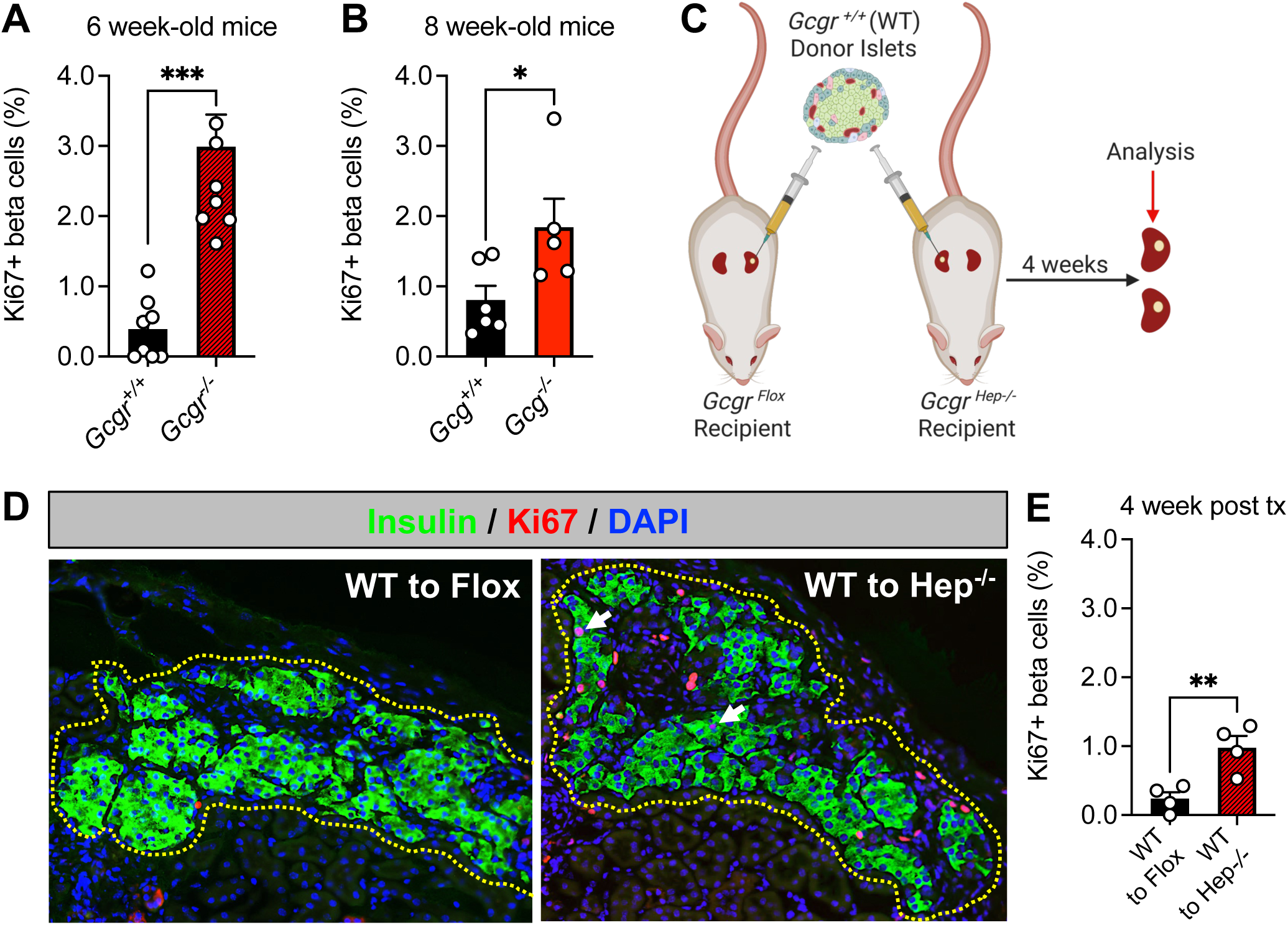
Genetic interruption of glucagon signaling stimulates beta cell proliferation in pancreatic and transplanted mouse islets. (**A-B**) Quantification of pancreatic islet beta cell proliferation in (**A**) 6 week-old *Gcgr^+/+^* (black bar, all males) and *Gcgr^-/-^* (red striped bar, all males) and (**B**) 8 week-old *Gcg^+/+^* (black bar) and *Gcg^-/-^* (red bar) mice (n=2-4 females and 3 males per group, unpaired t test, ***p < 0.001 versus *Gcgr^+/+^*, *p < 0.05 versus *Gcg^+/+^*). (**C**) Schematic of approach for subcapsular renal transplantation of *Gcgr^+/+^* (wild type, WT) donor islets into control (*Gcgr^Flox^*) or liver-specific *Gcgr* knockout (*Gcgr^Hep-/-^*) recipient mice. Created with BioRender.com (**D**) Representative images of islet grafts from WT to Flox and WT to Hep^-/-^ recipients after four weeks. Grafts are immunostained for insulin (green), Ki67 (red) and DAPI (blue). White arrows indicate Ki67+ insulin+ cells. Dashed yellow lines indicate kidney-graft boundary. (**E**) Quantification of beta cell proliferation in transplanted islets from WT to Flox (black bar) and WT to Hep^-/-^ (red striped bar) groups (n=4 males per group, unpaired t test, **p < 0.05 versus WT to Flox).

**Supplemental Figure 2.**
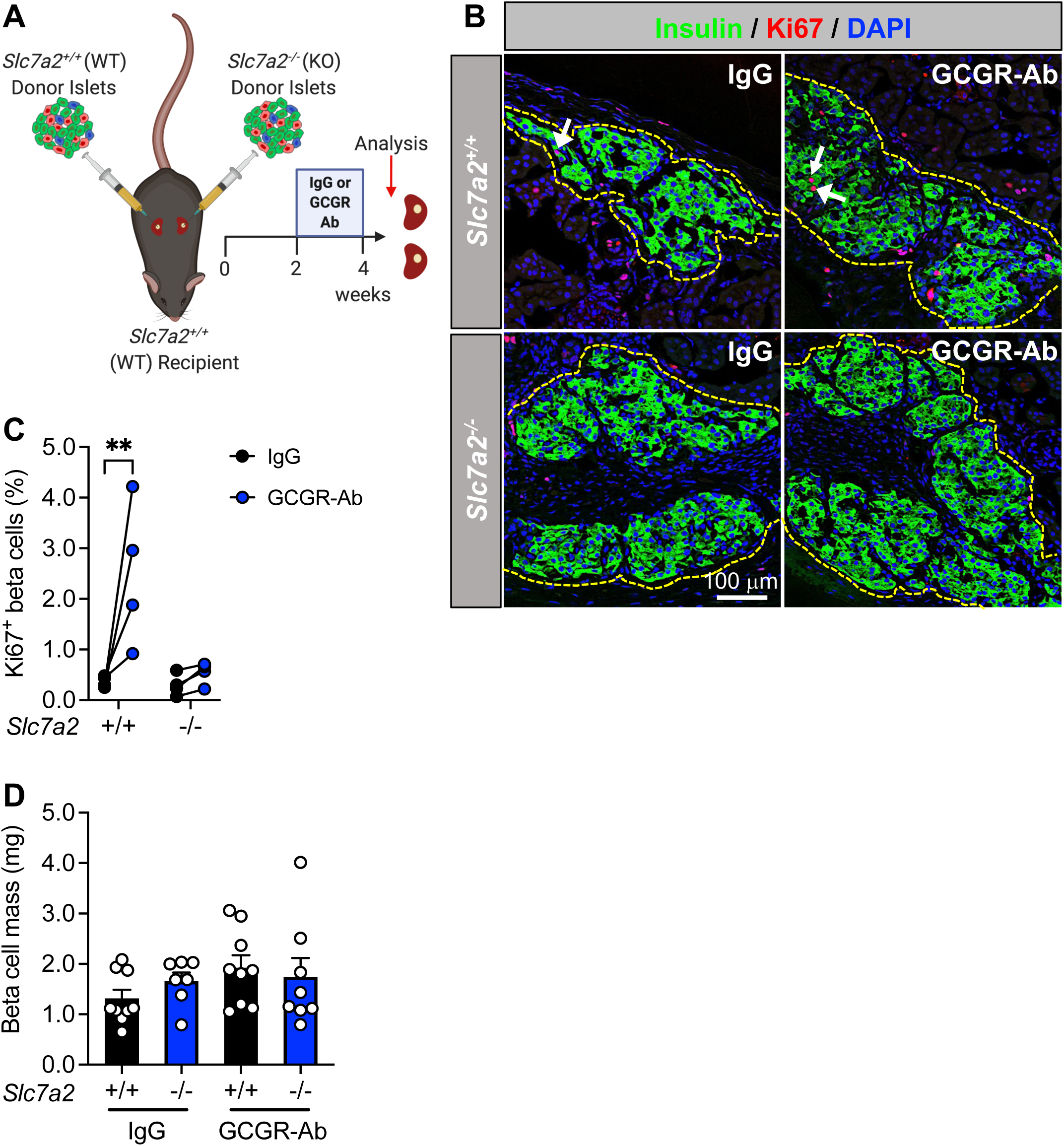
SLC7A2-dependent stimulated beta cell proliferation is islet autonomous. (**A**) Schematic of approach for subcapsular renal transplantation of *Slc7a2^+/+^*(wild type, WT) and *Slc7a2^-/-^* (KO) donor islets into *Slc7a2^+/+^* (WT) recipient mice followed by control IgG or GCGR-Ab treatment. Created with BioRender.com (**B**) Representative images of *Slc7a2^+/+^* (upper row) and *Slc7a2^-/-^* (bottom row) islet grafts from *Slc7a2^+/+^* kidney capsules after two weeks of IgG or GCGR-Ab treatment. Grafts are immunostained for insulin (green), Ki67 (red) and DAPI (blue). White arrows indicate Ki67+ insulin+ cells. Dashed yellow lines indicate kidney-graft boundary. (**C**) Quantification of beta cell proliferation in transplanted islets from *Slc7a2^+/+^*and *Slc7a2^-/-^* donors treated with IgG (black circles) or GCGR-Ab (blue circles; n=2 females and 2 males per treatment group, two-way ANOVA with Fisher’s LSD test, **p < 0.01 versus IgG treated). (**D**) Quantification of pancreatic islet beta cell mass in *Slc7a2^+/+^* (black bars) and *Slc7a2^-/-^* (blue bars) IgG or GCGR-Ab-treated mice (n=2-5 females and 3-6 males per group).

**Supplemental Table 1.** Human Islet Donor Information.

| Donor ID | Age | Ethnicity/Race | Sex | BMI<br>(kg/m<sup>2</sup>) | HbA1c<br>(%) | Cause of<br>Death | Islet<br>Source |
| --- | --- | --- | --- | --- | --- | --- | --- |
| AELC213 | 10 | Hispanic/Latino | F | 25.4 | N/A | Head<br>Trauma/Blunt<br>Injury | Other |
| AFEA331 | 45 | Black | M | 29.3 | 5.0 | CVA/ stroke | IIDP |
| AIFV371 | 28 | Hispanic/Latino | F | 24.7 | 5.0 | CVA/ stroke | HPAP |
| 1 | 32 | N/A | M | 29.5 | N/A | N/A | IIDP |
| 2 | 47 | N/A | M | 22.3 | N/A | N/A | IIDP |
| 3 | 55 | N/A | M | 28.4 | N/A | N/A | IIDP |
| 4 | 43 | N/A | M | 29.6 | N/A | N/A | IIDP |
| 5 | 46 | N/A | M | 28.8 | N/A | N/A | IIDP |
| 6 | 41 | N/A | F | 31.1 | N/A | N/A | IIDP |
| 7 | 47 | N/A | F | 25.6 | N/A | N/A | IIDP |
| 8 | 52 | N/A | M | 33.2 | N/A | N/A | IIDP |

## Notes

### Competing Interest Statement

Hai Yan is an employee of REMD Biotherapeutics Inc.

